# Canavanine-based assay for gross chromosomal rearrangements reveals genome instability hotspots and modulating genes in fission yeast

**DOI:** 10.64898/2026.05.15.725498

**Authors:** Anissia Ait Saada, Cassandre Ollivier, Alex B. Costa, Kevin Moreau, Sarah A.E. Lambert, Kirill S. Lobachev

## Abstract

Gross chromosomal rearrangements are a hallmark of many diseases and cancers. The study of their biogenesis and the mechanisms underlying their formation is greatly facilitated by the availability of genetic reporter assays in model organisms. We present here a novel GCR assay developed in fission yeast, a highly relevant model for understanding genome instability related to human biology. The reporter employs canavanine counter-selection to detect GCRs within a chromosomal context. Using this assay, we identified natural hotspots for GCRs, including inverted long terminal repeats (IR-LTRs). Structural analysis of GCR events showed that IR-LTR–induced GCRs mainly result in either terminal deletions with adjacent inverted duplications or repair via long-range break-induced replication (BIR). Deleting IR-LTRs reduces the GCR rate and reveals another hotspot driven by BIR between homeologous aldo/keto reductase genes on opposite arms of chromosome I. This is the first evidence that BIR can occur in *S. pombe* on long tracks reaching up to 600 kb. Besides highlighting genome rearrangement hotspots, the assay also identifies regulators of genome instability in fission yeast. Loss of Nup132, a component of the nuclear pore complex, increases IR-LTRs-induced GCRs, while the budding yeast homolog Nup133 has no effect on the stability of a structurally similar IR. In contrast, disrupting *djc9*, which encodes a conserved histone H3–H4 binding protein, decreases GCR rates. Overall, this sensitive GCR assay enables the identification of factors that control spontaneous and fragile motif–induced chromosomal instability, including those conserved in humans but lost through evolution in other organisms.

## Introduction

Genome instability is a hallmark of many diseases and plays a key role in the development and progression of tumors. Large-scale genome changes, including translocations, large deletions, inversions, and amplifications—referred to as gross chromosomal rearrangements (GCRs)—are the main type of genomic instability in numerous cancers (Al-Rawi and Bakhoum, 2022; Drews et al., 2022; Schuy et al., 2022).

Genetic assays, especially in model organisms, are essential for providing quantitative data on mutations and chromosomal rearrangements, helping to uncover the underlying molecular mechanisms of their biogenesis. The effectiveness of these assays depends on developing genetic markers that allow for accurate tracking and measurement of mutational events. Our current understanding of GCR formation mainly comes from studies in the yeast *S. cerevisiae*, where a GCR assay was developed in R. Kolodner’s laboratory (Chen and Kolodner, 1999). This assay relies on using two counter-selectable markers: *CAN1* and *URA3*, which confer resistance to canavanine and 5-FOA, respectively, when those genes are lost in the nonessential part of chromosome V (Putnam and Kolodner, 2010). A modification of this GCR assay involves the use of *CAN1* and *ADE2*, where the loss of both markers results in canavanine-resistant, red-colored colonies (Ait Saada et al., 2021a; Narayanan et al., 2006). The GCR cassette is inserted on the left arm of chromosome V, downstream of the essential PCM1 gene. GCRs are manifested as terminal chromosomal deletions accompanied by *de novo* telomere additions, interstitial deletions, or more complex rearrangements such as translocations or fold-back inversions (Li et al., 2020; Putnam and Kolodner, 2017). The spontaneous level of GCRs is usually extremely low (∼3x10^-8 to 1x 10^-9 event/cell/division), which provides a background instrumental in the identification of yeast genome instability suppressing (GIS) genes (Chen and Kolodner, 1999; Putnam et al., 2016, 2009; Putnam and Kolodner, 2017; Srivatsan et al., 2019), where mutations in their human homologues are enriched in several human cancers. The rate of spontaneous GCRs and their structure are highly influenced by the nature of the sequence between the GCR cassette and the first essential gene. For example, insertion of Ty1 retrotransposon led to a 380-fold increase in GCR rates in wild-type strains, where a major class of rearrangements is represented by nonreciprocal translocation events mediated by recombination between genome-wide dispersed Ty elements (Chan and Kolodner, 2011).

Despite key discoveries in *S. cerevisiae* regarding GCR etiology, our understanding of both GIS genes and genome instability-promoting (GIP) genes in eukaryotes remains incomplete. In addition, 356 genes are conserved between the fission yeast *S. pombe* and humans but are absent in budding yeast, and which of them are involved in the maintenance of genome stability remains to be assessed. Therefore, developing a convenient assay to systematically screen how the deficiency of *S. pombe*-specific genes—many of which are orthologs of disease-causing human genes—affects GCRs is necessary. So far, one study has reported the use of a GCR assay in fission yeast in a chromosomal context (Irie et al., 2019) and several other studies have used extra-chromosomes to score for rearrangements derived from chromosome 3 to study centromere-mediated GCRs (Irie et al., 2019; Nakamura et al., 2008a; Okita et al., 2019; Onaka et al., 2020; Pai et al., 2023). The assay developed by Irie et al. uses the *ura4* and *tk* genes for counterselection on 5-FOA and FuDr (Irie et al., 2019), while the second employs an extra mini-chromosome (derived from chromosome 3) containing a GCR cassette. These assays demonstrated a novel role for the shelterin components Taz1 and Rap1, as well as heterochromatin, in suppressing GCRs. However, they have some drawbacks, including the requirement to work in specific genetic backgrounds to incorporate the nucleotide analog, the use of the *ura4* marker, widely employed in fission yeast genetic manipulations, or the investigation of GCRs outside the normal chromosomal context. Surprisingly, there is no reliable canavanine-based genetic assay in *S. pombe* to measure genome instability, although basic amino acid/canavanine uptake is well documented (Aspuria et al., 2008). The challenge in creating a dependable genetic assay using canavanine counterselection lies in two issues. First, the loss of function of a single arginine permease in *S. pombe* is not enough to induce strict canavanine resistance (Ait Saada et al., 2022; Aspuria and Tamanoi, 2008). Second, the *can1-1* allele, which is the only single mutation capable of conferring canavanine resistance, is a dominant gain-of-function mutation (Ait Saada et al., 2022; Yang et al., 2022).

Here, to develop a versatile chromosomal GCR assay in *S. pombe*, we used our knowledge of canavanine uptake and the requirement for a multi-marker cassette in GCR assays. The assay is based on a cassette inserted into a non-essential region of the left arm of chromosome I, consisting of two arginine permease genes, *cat1* and *aat1*, and the colorimetric marker *ade7*. GCR rates in fission yeast are several orders of magnitude higher than those in the original budding yeast GCR assay; this difference is due to the presence of repetitive elements in the non-essential part of the left arm of chromosome I. Using this assay, we identified natural hotspots for GCR formation, including a region with two long terminal repeats (IR-LTRs) that are 211 bp apart and share over 96% homology in 176 bp of the inverted repeats. We show that GCRs induced by IR-LTRs follow a specific pattern characterized by terminal deletions coupled with adjacent inverted duplications. Often, broken ends are repaired by long-range break-induced replication (BIR), leading to nonreciprocal translocation. Deleting IR-LTRs results in a threefold decrease in GCR rate and reveals another GCR hotspot promoted by BIR involving recombination between two homeologous aldo/keto reductase genes located on different arms of chromosome I. Additionally, this assay can be used to identify and characterize genes that suppress or promote genome instability in fission yeast. We found that deleting *nup132*, which encodes a component of the nuclear pore complex, destabilizes IR-LTRs, whereas disrupting the budding yeast homolog Nup133 does not affect the fragility of similarly structured IRs. Nup132 deficiency also induces GCRs in strains lacking IR-LTRs. Conversely, disrupting *djc9*, a gene encoding a histone H3-H4 binding protein conserved in humans but not in budding yeast, decreases the GCR rate in strains with or without IR-LTRs. This study demonstrates that the newly developed sensitive GCR assay provides an opportunity to identify novel factors governing spontaneous and fragile motif-induced chromosomal instability.

## Results

### Canavanine-resistance-based GCR assay in fission yeast

Terminal non-essential regions in *S. pombe* have been identified (Tashiro et al., 2017). Here, we inserted the GCR cassette approximately 50 kb away from the telomere and 50 kb downstream of the first essential gene, *pdc101*, on the left arm of ChrI (I:50132-51288). Although the GCR cassette location is mapped roughly 50 kb from the end of chromosome I in the Pombe database (https://www.pombase.org), it is about 30 kb farther from the telomere, according to Tashiro et al., where unsequenced regions are included (Tashiro et al., 2017). For simplicity, the coordinates from the Pombe database are shown. As shown in Figure 1A, the CGR cassette includes the *aat1, cat1*, and *ade7* genes. The double *aat1Δ cat1Δ* mutant exhibits a synergistic increase in canavanine resistance compared to each single mutant on ammonium chloride-containing medium (Aspuria and Tamanoi, 2008) (Figure S1). Incorporating both genes into the GCR cassette allows us to use a stringent medium for canavanine selection, eliminating background growth. In fact, using low levels of ammonium chloride (0.5 g/L instead of 5 g/L) reduces background growth in wild-type cells without affecting the growth of canavanine-resistant cells (Ait Saada et al., 2022). The third genetic marker, *ade7*, affects the adenine biosynthesis pathway. When *ade7* is mutated or lost, cells accumulate a red pigment due to the buildup of the intermediate phosphoribosylaminoimidazole carboxylate (Fisher, 1969). Loss of the GCR cassette can be selected on media supplemented with canavanine and low levels of adenine. Therefore, the GCR rates can be determined by counting canavanine-resistant red colonies (indicating loss of *aat1, cat1*, and *ade7*) (Figure 1).

**Figure 1.**
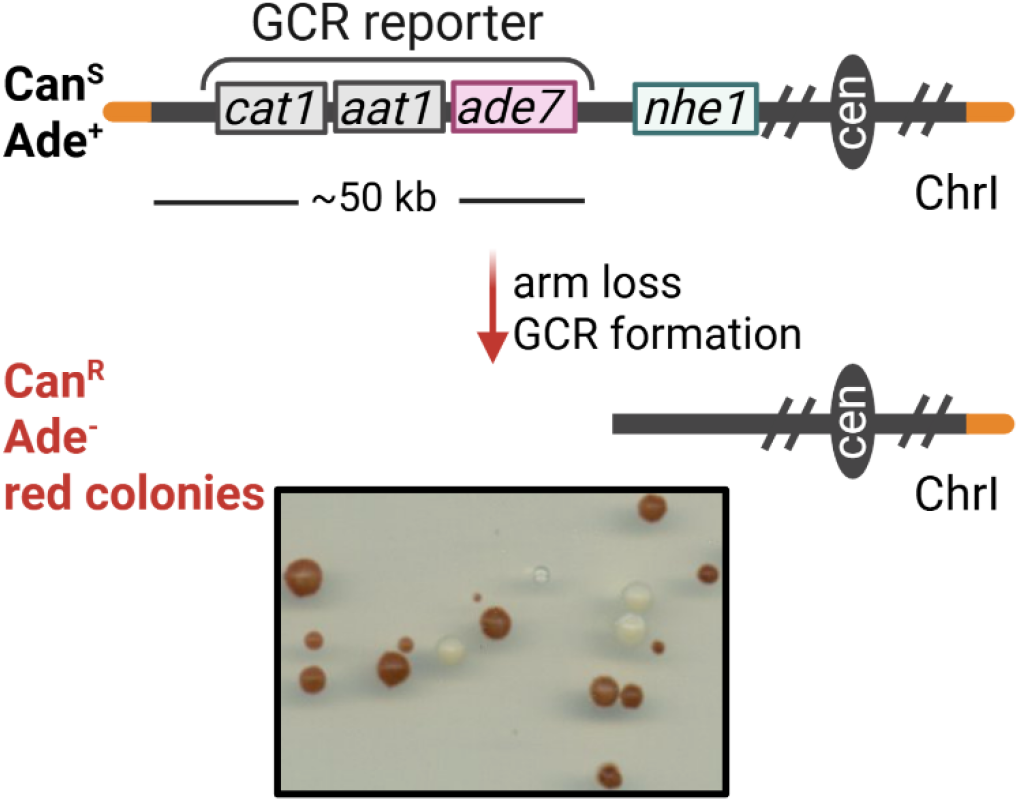
Overview of the canavanine-based GCR assay in *S. pombe*. The GCR cassette, composed of *cat1, aat1*, and *ade7* genes, is inserted on the non-essential left arm of chromosome I, ∼ 50 kb away from the telomere. The genes composing the cassette were deleted from their endogenous location by *delitto perfetto* (Fréon et al., 2023). Cells expressing the cassette are canavanine sensitive, while the loss of the cassette renders cells canavanine-resistant and red (CanR Ade-). An example of red canavanine-resistant colonies, corresponding to cells harboring GCR events, is shown.

### Analysis of GCR events reveals a recurrent breakpoint at the inverted retroelements

The rate of spontaneous GCRs in WT cells was unexpectedly high, corresponding to 1.39x10^-5^ event/cell/division (**Table 1**). This rate is several orders of magnitude higher than the rates determined for i) the right arm of chromosome I in fission yeast (Irie et al., 2019) or for the left arm of chromosome V in budding yeast (Chen and Kolodner, 1999; Narayanan et al., 2006).

**Table 1.**
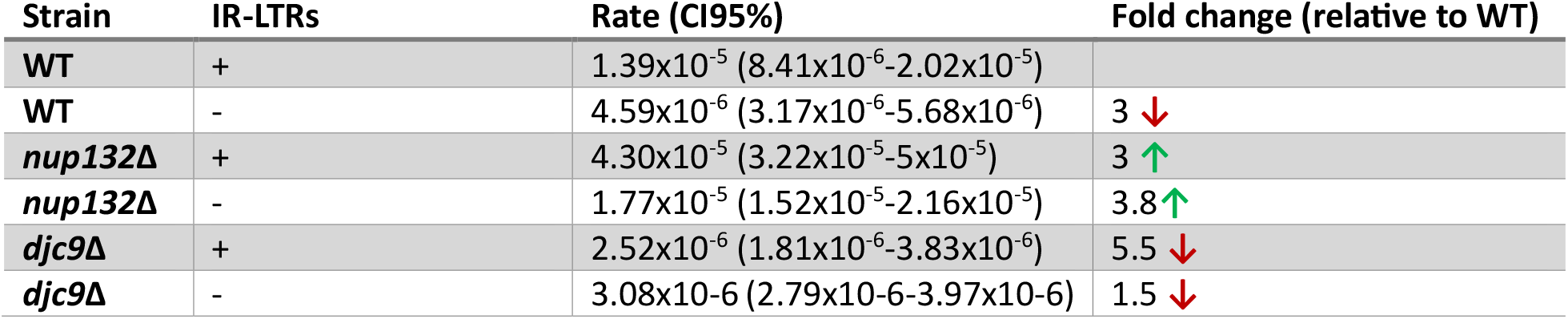
GCR rates in wild-type, *nup132Δ* and *djc9Δ* mutants with and without IR-LTRs.

| Strain | IR-LTRs | Rate (CI95%) | Fold change (relative to WT) |
| --- | --- | --- | --- |
| WT | + | $1.39 \times 10^{-5}$ ($8.41 \times 10^{-6}$-$2.02 \times 10^{-5}$) | |
| WT | - | $4.59 \times 10^{-6}$ ($3.17 \times 10^{-6}$-$5.68 \times 10^{-6}$) | 3 ↓ |
| *nup132Δ* | + | $4.30 \times 10^{-5}$ ($3.22 \times 10^{-5}$-$5 \times 10^{-5}$) | 3 ↑ |
| *nup132Δ* | - | $1.77 \times 10^{-5}$ ($1.52 \times 10^{-5}$-$2.16 \times 10^{-5}$) | 3.8 ↑ |
| *djc9Δ* | + | $2.52 \times 10^{-6}$ ($1.81 \times 10^{-6}$-$3.83 \times 10^{-6}$) | 5.5 ↓ |
| *djc9Δ* | - | $3.08 \times 10^{-6}$ ($2.79 \times 10^{-6}$-$3.97 \times 10^{-6}$) | 1.5 ↓ |

To characterize GCR events, we performed whole-genome sequencing on 13 Can^R^ Ade^-^ clones. In all of them, the phenotype resulted from the loss of the region containing the GCR cassette, as all clones showed a terminal deletion of the left arm of chromosome I (Figure 2A). The deletion was found to be upstream of the GCR cassette, but none of the clones exhibited a deletion beyond the *pcd101* essential gene (Figure 2A). In most GCR clones, the breakpoint appeared to be located within a 400 bp window (between 98.6 and 99 kb from the telomere) that contains two retroelements, *SPLTRA*.*11* and SPLTRA.*12*, in inverted orientation. One GCR clone (Clone C7) showed a breakpoint 2kb away from the location of these retroelements, at 101.3 kb (Figure 2A).

**Figure 2.**
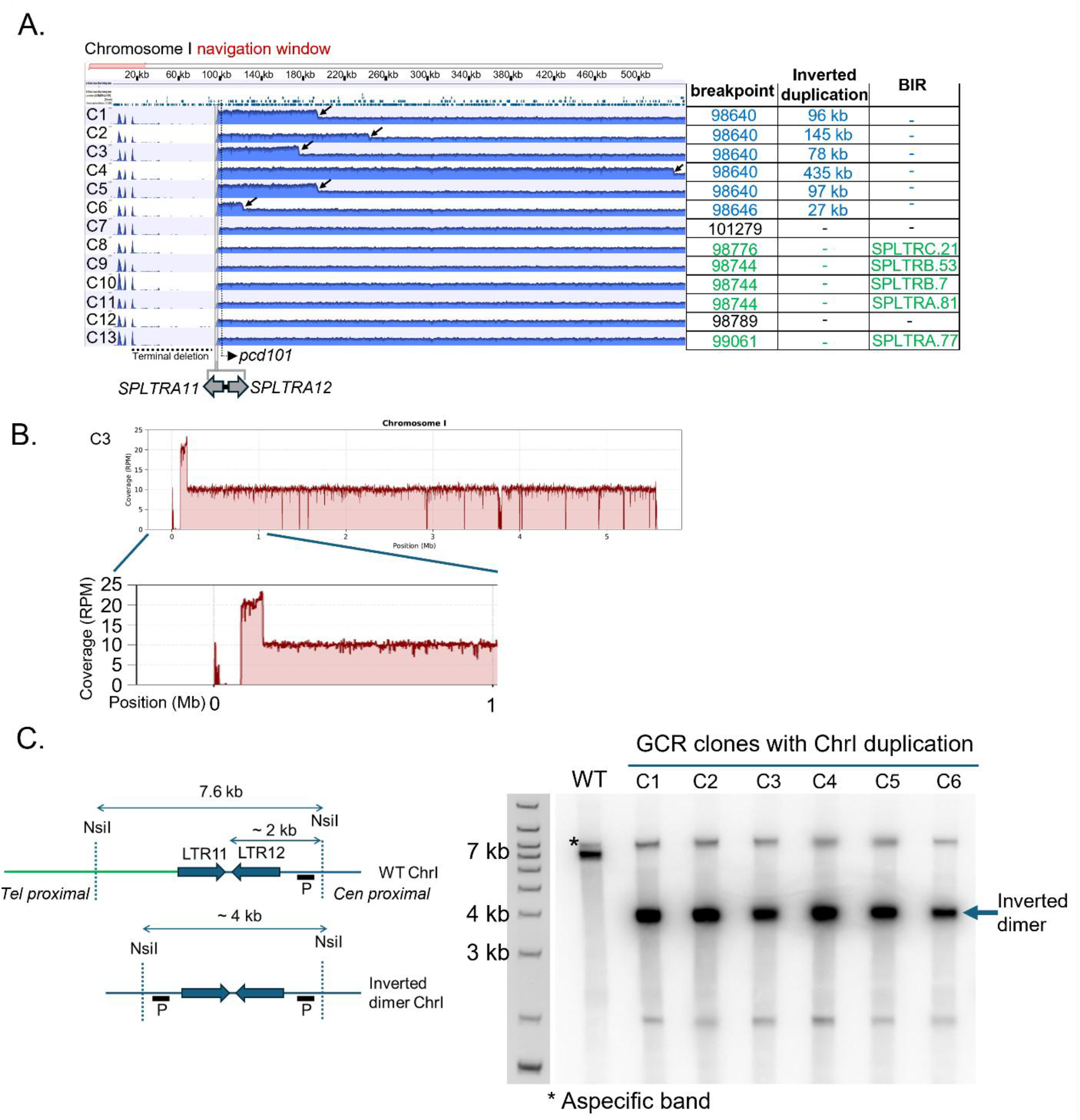
Structural analysis of GCR events in WT cells by Illumina sequencing reveals different classes of rearrangements. **(A)** Sequencing analysis of 12 GCR clones. Left panel: overview of the terminal left arm of chromosome I. A window encompassing 600 kb of the terminal left arm of chromosome I is shown. All clones harbor a terminal deletion manifested by the absence of mapped reads in the 100 kb of chromosome I. The black arrows indicate the end of the adjacent duplication in the concerned clones. Right panel: categorization of the GCR events according to their breakpoint, the presence and length of an adjacent duplication and BIR event. **(B)** and **(C)** Read depth analysis of GCR clones for the indicated chromosome. **(B)** Read depth analysis of a GCR clone showing adjacent duplication. **(C)** The adjacent duplication detected in GCR events corresponds to an inverted duplication. Analysis of inverted duplication by Southern blot. Left panel: scheme of the left arm of chromosome I and the inverted duplication position. The NsiI restriction sites, the size of the DNA fragments and the location of the probe are indicated. Right panel: detection of inverted duplication in GCR clones by Southern blot. Genomic DNA was submitted to NsiI digestion. WT cells exhibit the ∼ 7.6 kb band while the GCR clones exhibit a ∼ 4 kb size band expected in the presence of an inverted duplication at the *SPLTRA11*-*SPLTR12* locus.

Although the breakpoints are in the same region, different GCR classes can be distinguished. First, a subset of GCR clones (6/13, Clones C1-C6 in blue on Figure 2A) showed a duplication next to the breakpoint on the remaining left arm of chromosome I (see Figure 2B for an example). The size of this duplication varied among the clones, ranging from 24 to 435 kb (Figure 2A, black arrows indicate the end of the duplication and the size of the duplication is indicated on the right panel). The two naturally occurring retroelements, *SPLTRA*.*11* and *SPLTRA*.*12*, are arranged in an inverted orientation and form a quasi-palindrome consisting of 96% homologous 176 bp repeats with a 211 bp spacer (hereafter called IR-LTRs). DNA inverted repeats are hotspots for genome instability, and the extent of this instability depends on the homology of the arms and the length of the spacer (Ait Saada et al., 2023; Lobachev et al., 2000; Ramakrishnan et al., 2018). It has been shown that inverted repeats separated by several kilobases or exhibiting up to 86% homology can induce chromosome instability (Lobachev et al., 1998). Inverted repeats can cause chromosome rearrangements through the formation of a palindromic dicentric chromosome, and breakage of this structure can produce a chromosome with an inverted duplication (Narayanan et al., 2006). Therefore, we examined whether the duplication near the breakpoint in six clones is an inverted dimer. To do this, we analyzed the structure of the duplications in these GCR clones using Southern blot analysis of restriction fragments (Figure 2C). In the wild-type strain, only a 7.6 kb fragment was observed, but all GCR clones analyzed showed a band around 4 kb, consistent with the expected size for an inverted dimer formation (Figure 3C). This indicates that in this first class of GCRs, arm loss involves a foldback inversion mechanism mediated by the IR-LTRs, resulting in a dicentric palindromic chromosome and resolution of which leads to a chromosome with an inverted duplication. The presence of the quasi-palindrome at the breakpoint suggests that its inherent features—such as being prone to breakage or fold-back formation in single-strand DNA—may trigger GCRs. To understand how the broken dicentric chromosome healed, we identified the unmapped reads at the centromere-proximal side of the duplication. Of the six clones exhibiting inverted duplication, the broken inverted dimer chromosome was stabilized by acquiring a telomere in five cases (see Figure S2A for examples). The remaining clone showed a more complex rearrangement, consisting of an additional second distal duplication (from 1.64 Mb to 2.37 Mb) and a deletion of the right arm of chromosome I (Fig S2B, red arrow). The left side of the duplication showed an unmapped sequence (represented by a black circle in Fig S2B) corresponding to the end of the initial duplication (i.e., at position 535 kb), consistent with the repair of the broken chromosome by break-induced replication (invasion of the broken dicentric chromosome of the region and synthesis). Surprisingly, the unmapped sequences on the right side of the distal duplication (represented by a red circle in Fig S2B) did not correspond to telomeric sequences but rather to the sequence adjacent to the breakpoint on the right arm of chromosome I. These data suggest that after or during dicentric chromosome breakage, a secondary break occurred at the right terminal region of chromosome I. Each broken end subsequently invaded the same chromosomal interval, spanning 1.64 Mb to 2.37 Mb: the initial break invaded at 1.64 Mb, whereas the secondary right-terminal break invaded at 2.37 Mb. This indicates that the resulting rearrangement is likely a circular chromosome I.

**Figure 3.**
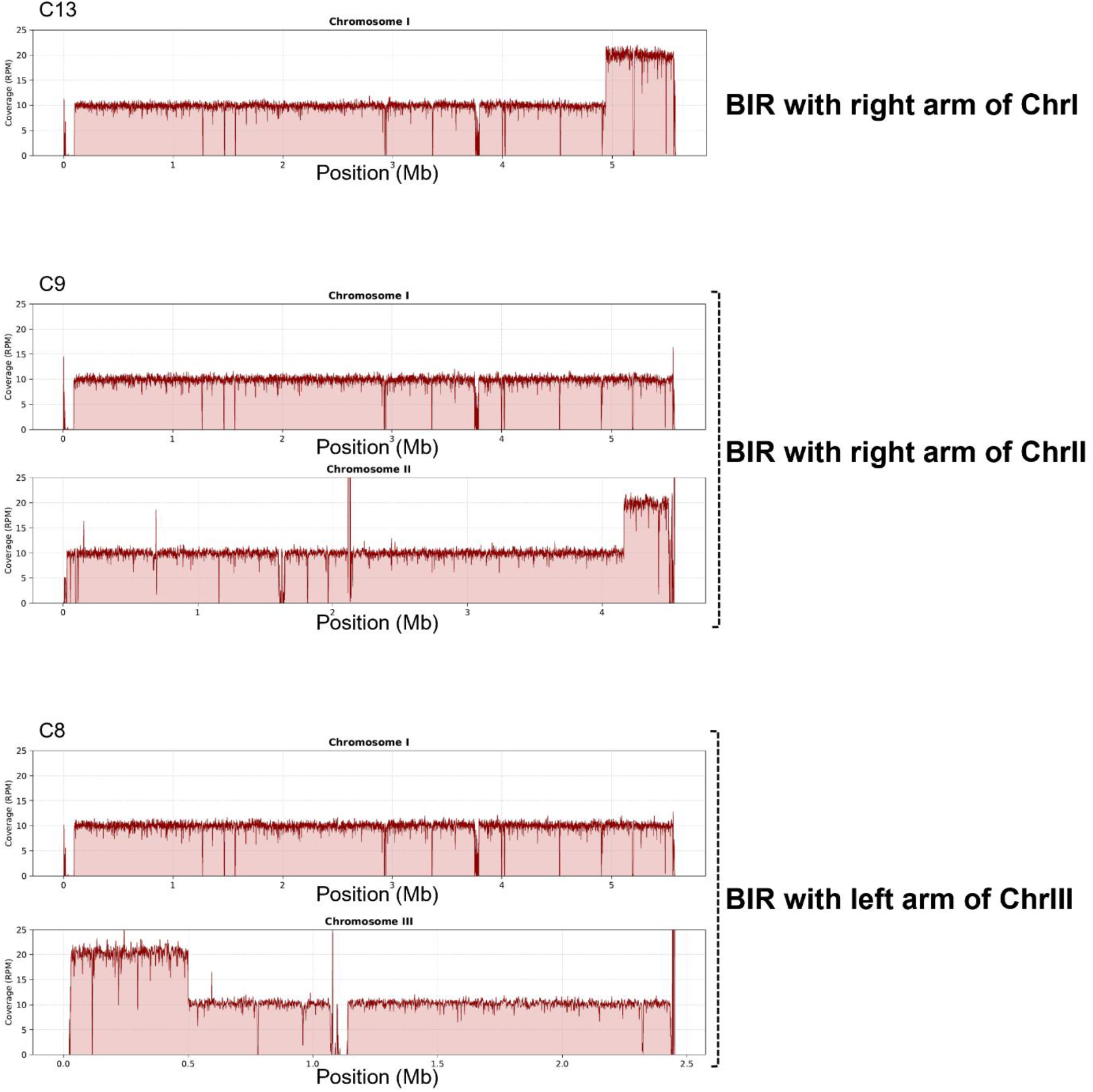
Structural analysis of GCR events in WT reveals that a subset of rearrangements occurs via BIR. Examples of BIR events illustrated by the detection of an ectopic duplication on the right arm of chromosome I (top panel) or chromosome II (middle panel) or the left arm of chromosome III (bottom panel).

The second class of GCRs comprises five out of the 13 examined clones, displaying a terminal deletion accompanied by the invasion of another retroelement located elsewhere in the genome, representing a break-induced replication (BIR) event (Figure 2A, clones in green on the right panel). This is supported by the detection of sequences at the breakpoint corresponding to the invaded region. The invaded retroelements were situated either on the right arms of chromosome I, the left or right arm of chromosome II, or the left arm of chromosome III, and BIR tracts ranged from 63 to 636 kb (Figure 3). Interestingly, each of the four BIR-like events involved a different retroelement (*SPLTRA*.*77, SPLTRA*.*81, SPLTRB*.*7, SPTLRB*.*53*, and *SPLTR20*). Lastly, the third class of rearrangements includes terminal deletions (2/13) without duplication or BIR events (Figure 2A, clones C7 and C12), resulting from telomere addition since the unmapped sequences at the breakpoints harbor the telomeric sequence repeat GGTTAC (see Figure S2C for examples).

### Deletion of *SPLTRA*.*11* and *SPLTRA*.*12* leads to a decrease in GCR rates and reveals another hotspot for rearrangements

Repetitive elements are known to influence GCR levels in budding yeast. For instance, inserting a single Ty1 element into a non-essential terminal region of chromosome V increased GCR rates by approximately 380-fold in a wild-type strain (Chan and Kolodner, 2011), while introducing a quasi-palindrome containing 94% homologous inverted repeats separated by a 200 bp spacer raised GCR rates by a factor of 1,000 (Ait Saada et al., 2023). Since the breakpoints occur within a quasi-palindrome composed of two repetitive elements, we tested whether the high GCR rate is attributable to the inverted repeats. Deleting IR-LTRs reduces the GCR rate by 3-fold (Table 1), indicating that they trigger rearrangements but are not the only factors. This suggests there may be other, possibly less potent, hotspots for GCR formation.

Eleven Can^R^Ade^-^ clones were sequenced to map the rearrangement breakpoints. As expected, the breakpoints were mapped closer to the GCR cassette insertion site compared to the GCR clones derived from the strain harboring the IR-LTRs (Figure 4A). Additionally, the position of the breakpoint was more variable. While all clones with IR-LTRs exhibited approximately 98 kb deletions, those without IR-LTRs showed the following five breakpoint positions: 55 kb (2/11, clones C6 and C7), 57 kb (1/11, clone C8), 60 kb (5/11, clones C1 to C5), 68 kb (2/11, clones C9 and C10), and 97 kb (1/11, clone C11) away from the telomere. As expected, none of the GCR events displayed a foldback inversion in the absence of IR-LTRs since no additional reads were mapped adjacent to the breakpoint (Figure 4, left panel), as compared to WT cells (Figure 2A, left panel clones C1 to C6). Several classes of rearrangements were identified. In the first one (5/11, C1 to C5 in green on Figure 4A), where the breakpoint is located approximately 60 kb from the telomere, this breakpoint falls within the NADP-dependent aldo/keto reductase gene (SPAC977.14C), and is accompanied a duplication of the right arm of chromosome I at the location of another NADP-dependent aldo/keto reductase gene (SPAC750.01) located ∼ 23 kb away from the telomere (Figure 4B). SPAC977.14C (left arm of ChrI) and SPAC750.01 (right arm of ChrI) are paralogs that share 96% homology. Therefore, this first class of GCRs corresponds to a BIR event involving the two NADP-dependent aldo/keto reductase genes. Surprisingly, only clone C1 showed a clear terminal deletion, whereas in the others (clones C2 to C5), the deletion seems partial, given the presence of the first 35 kb DNA sequence (Figure 4A, left panel). This indicates that the initial BIR involving two NADP-dependent aldo/keto reductase genes switched to the left arm of the broken molecule to complete chromosome repair. In budding yeast, it has been shown that BIR is prone to template switching (Stafa et al., 2014), which can explain this class of GCRs. The second class (in black on Figure 4A) corresponds to telomere addition events, as no secondary rearrangement was detected and clear telomeric sequences were identified at the breakpoints for two clones (3/11, clones C8, C9, and C10). The third class (2/11), not observed in WT cells, consisted of interstitial deletions involving retrotransposon elements *SPLTRA*.*7* and *SPLTRA*.*1* in one case and *SPLTRA*.*7* and *SPLTRA*.*5* in another (Clones C6 and C7, in blue on Figure 4A). Finally, one clone (clone C11 on Figure 4A) exhibited a more complex rearrangement, consisting of the deletion of the left arms of chromosomes I and II (96 kb and 80 kb, respectively) and a duplication of a portion of chromosome I spanning from 2.52 Mb to 2.65 Mb (Figure 4C). None of the breakpoints were associated with telomeric sequences; instead, they mapped to a terminal sequence of chromosome II, corresponding to a region labeled as telomeric gap region. In addition, both the borders of the duplicated region ended with telomeric sequences. Therefore, the final GCR formed remains only partially understood.

**Figure 4.**
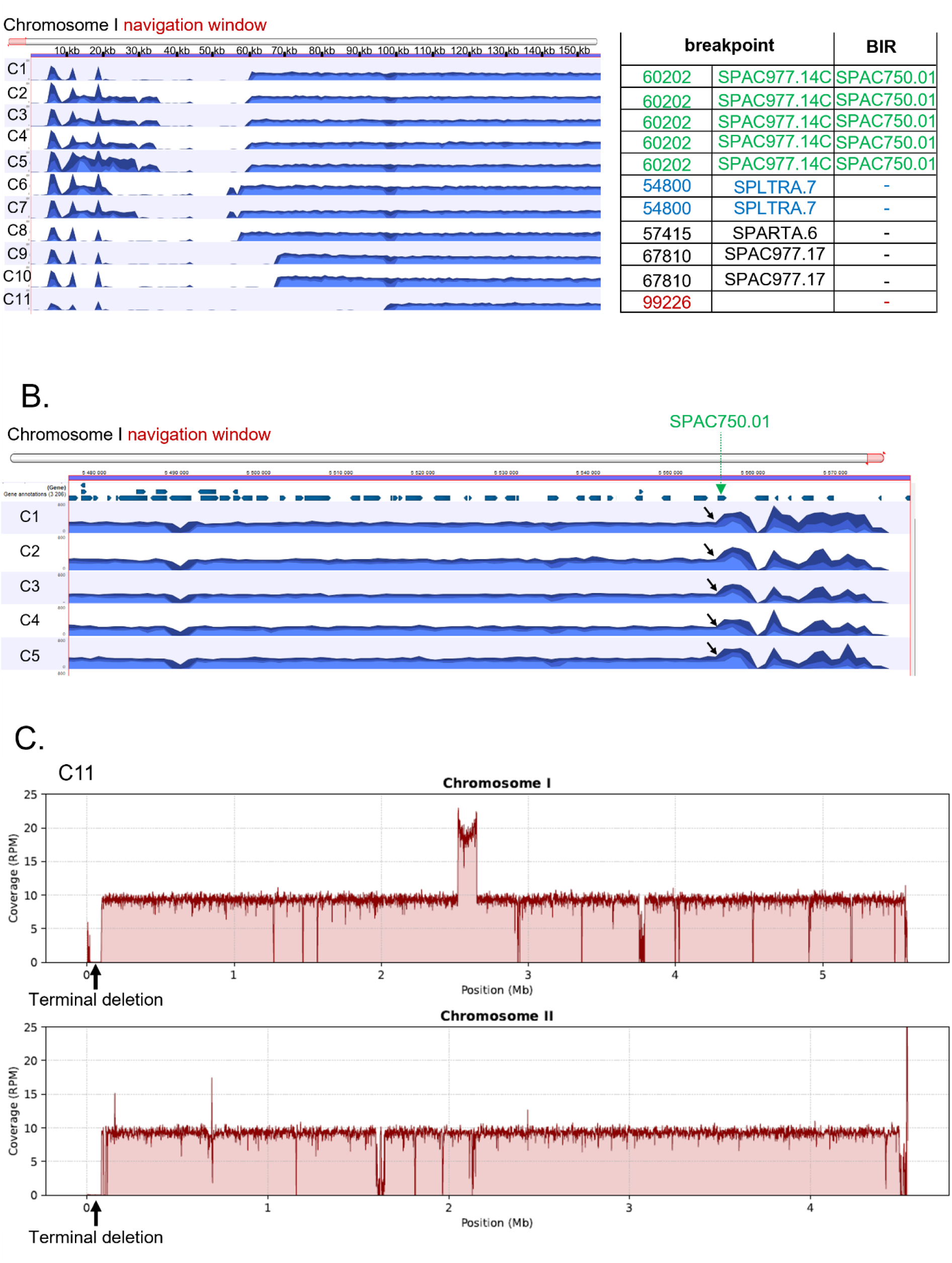
Structural analysis of GCR events in strains devoid of *SPLTRA11* and *SPLTR12* by Illumina sequencing. **(A)** Sequencing analysis of 12 GCR clones. Left panel: overview of the terminal left arm of chromosome I. Right panel: categorization of the GCR events according to their breakpoint and BIR event. **(B)** Example of homeologous recombination between *SPAC977*.*14C* (at the breakpoint) and *SPAC750*.*01* (right arm of chromosome I). This event is exemplified by the detection of an ectopic duplication on the right arm of chromosome I (black arrow) at the position of *SPAC750*.*01* (green arrow). **(C)** Example of secondary rearrangement in one GCR clone showing a chromosome I duplication and a terminal deletion of the left arm of chromosome II (black arrows).

These results demonstrate that the numerous repeat regions along the *S. pombe* chromosome significantly affect GCR recovery and determine rearrangement outcomes.

### Use of the GCR assay to identify GIS and GIP genes as hyper GCR and hypo GCR mutants

GCR assays in yeasts have been instrumental in identifying factors involved in maintaining genome stability. To confirm the robustness of the GCR assay in *S. pombe*, we used it to quantify genome rearrangements in mutants with established or uncharacterized roles in GCR formation.

We initially targeted a component of the nuclear pore complex within the Y-complex, the nucleoporin Nup132 (Holzer and Antonin, 2019). In all eukaryotes, a central component of the Y-complex is nucleoporin Nup133 (reviewed in (Holzer and Antonin, 2019)). Unlike in *S. cerevisiae* and humans, fission yeast has two nucleoporin isoforms, Nup131 and Nup132. While Nup131 is located in the cytoplasmic ring, Nup132 resides in the nucleoplasmic part of the Y-complex. Deletion of *NUP133* in budding yeast was found to increase the rate of spontaneous GCRs but not the rate of inverted repeat-induced GCRs (Putnam et al., 2012; Zhang et al., 2013) and Table S1). We found that its deletion increases the GCR rate by approximately 3-fold in both the presence and absence of IR-LTRs (Table 1). This indicates that the GCR assay can be used to identify hyper GCR mutants. In addition, our data suggests that while the role of the Y-complex in preventing genome instability manifested by GCRs appears conserved between budding and fission yeast, its role in suppressing inverted repeat-induced GCRs seems specific to fission yeast. Moreover, this suggests that the nuclear part of the Y-complex is essential for this function.

The other gene we targeted is *djc9*, a conserved gene in *S. pombe* and mammals. It encodes a histone H3-H4 binding protein that directs these histones for degradation (Ding et al., 2025). We found that deleting *djc9* results in a fivefold decrease in the GCR rate in the presence of IR-LTRs and a 1.5-fold reduction in their absence (Table 1). This suggests that limiting the availability of histone H3-H4 in the WT strains promotes the GCR formation in fission yeast.

## Discussion

We present a canavanine-based assay to score GCRs in fission yeast, thereby broadening the available tools for studying genome stability. To develop a reliable method for detecting GCRs in *S. pombe*, we engineered and optimized a GCR assay that relies on counterselection with canavanine and adenine auxotrophy. This assay allows for quantitative detection of genome rearrangements, and our results show that it consistently identifies rearrangement-prone genotypes and can differentiate between various types of genome instability phenotypes.

One advantage of the system we developed is the use of a single drug, canavanine, to counter-select for arm-loss events, combined with colorimetric selection for loss of the second marker. While canavanine is widely used in genetic assays in budding yeast, it remains less utilized in *S. pombe*. This difference arises because, in fission yeast, several redundant permeases, including Cat1 and Aat1, facilitate arginine uptake, whereas in *S. cerevisiae*, Can1 is the primary transporter. We capitalized on the need to use multiple markers to establish a GCR reporter that relies on canavanine-based counterselection. In line with this, Lyu et al. isolated four canavanine-resistant clones, all of which resulted from the loss of more than one amino acid transporter gene. Since most amino acid transporter genes are located at chromosome ends, two of the identified mutants involved diverse modes of chromosome II terminal deletions, where *aat1* is naturally located (Lyu et al., 2024). We chose not to use *ura4*, the most common counter-selectable marker in fission yeast, whose mutation confers resistance to 5-FOA, because it is also frequently used for positive selection and as a deletion marker for many genes.

The GCR cassette is positioned about 50 kb from the left arm of chromosome I and roughly 50 kb from the first essential gene, *pcd101*. Notably, this region contains multiple LTRs, enabling us to examine genome instability within the context of repetitive elements. This is particularly relevant since the human genome is rich in mobile and other repetitive elements (Smeds et al., 2025). As a result, the GCR rate in our system is several orders of magnitude higher than in other assays. For example, in Irie et al., spontaneous GCR rates were around 2×10^-9 events per cell per division. In that assay, the GCR cassette was inserted approximately 16.8 kb from the first essential gene, *sec16*, and GCRs mainly involved telomere additions (Irie et al., 2019). The flexibility of our system allows us to place the GCR cassette in any non-essential region of a chromosome, whether near repetitive elements or not.

Using this system, several types of rearrangements were detected and are recapitulated in Figure 5. In WT cells, most breakpoints occur in the region containing the *SPLTRA*.*11* and *SPLTRA*.*12* retroelements. Their inverted orientation makes this region a hotspot for genome instability, leading to the formation of GCRs (Figure 5A). In nearly half of the clones with IR-LTRs, GCRs are characterized by terminal deletion coupled with inverted duplication, with *SPLTRA*.*11* and *SPLTRA*.*12* retroelements at the center of symmetry. This indicates that dicentric formation and breakage are intermediate steps in the rearrangement process. IR-mediated dicentric chromosomes can form through different mechanisms: (i) the formation of secondary structures such as hairpins or cruciforms, nuclease attack, and breakage at IRs, which generates a hairpin-capped break (Ait Saada et al., 2021b; Narayanan et al., 2006), or (ii) fold-back at the homologous IR arms when the IR is within a single-strand DNA region, occurring when spontaneous or induced breakage and resection occur near the IRs (Ait Saada et al., 2023; Al-Zain et al., 2023; Deng et al., 2015; Li et al., 2020; Maringele and Lydall, 2004; VanHulle et al., 2007). We favor the first scenario because it is supported by several lines of evidence. First, nearly all the breakpoints (12 out of 13) identified in WT cells map to the *SPLTRA11-SPLTRA12*, despite the presence of other LTRs in this region (Figure 3). One breakpoint is only 2 kb from the IRs, likely indicating that breakage at *SPLTRA11-SPLTRA12* was followed by resection before DSB repair. Second, this finding aligns with previous results from Thomas Petes’ lab, which showed that inverted Ty retroelements act as natural fragile sites, promoting chromosomal breakage and mitotic recombination in budding yeast (Lemoine et al., 2008, 2005; Mieczkowski et al., 2006; St. Charles and Petes, 2013). Third, in budding yeast, repeats structurally similar to the fission yeast *SPLTRA11-SPLTRA12*, which are 94% identical and separated by 200 base pairs, strongly induce GCRs (Ait Saada et al., 2023).

**Figure 5.**
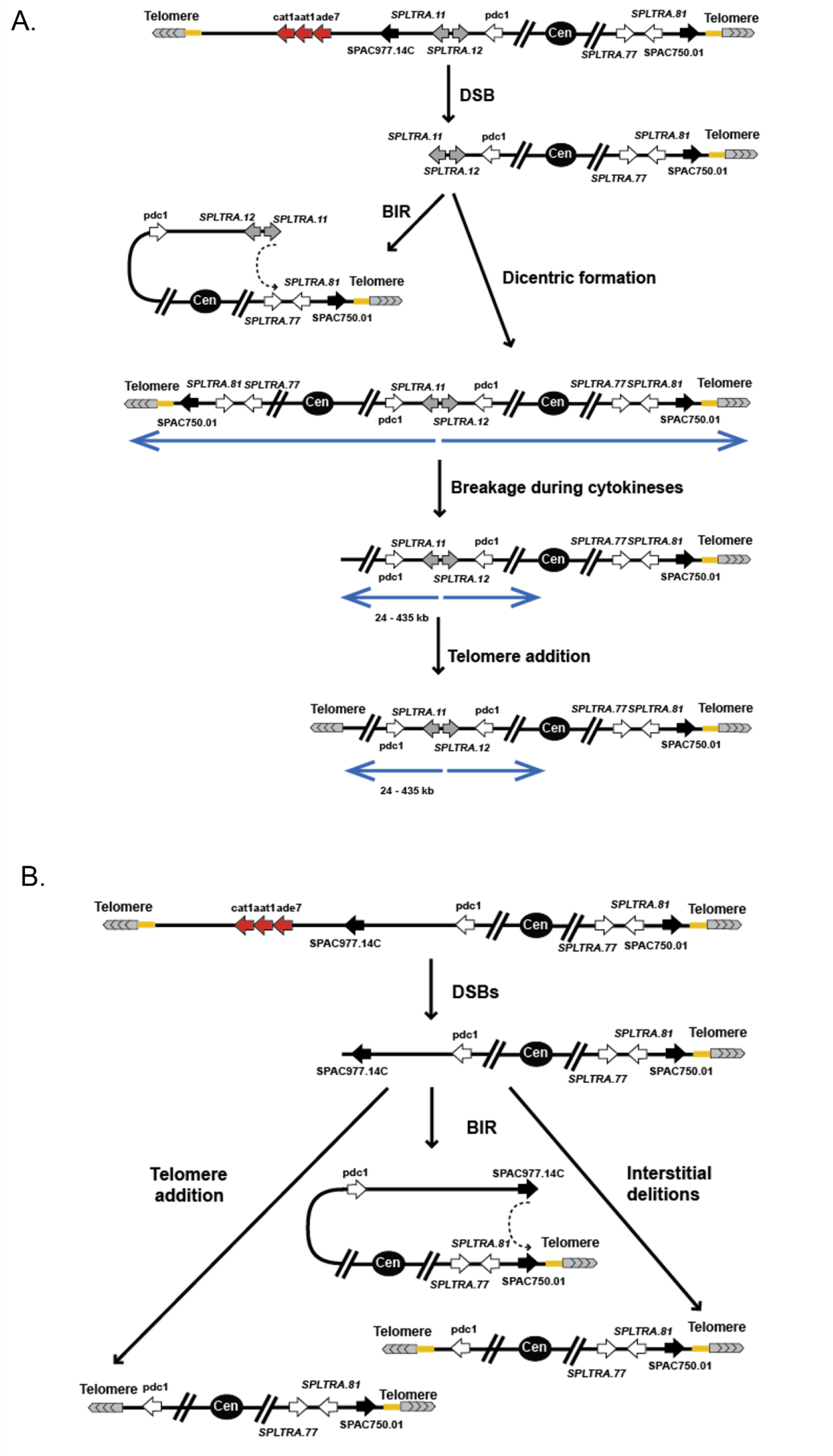
Mechanisms of GCR formation in fission yeast. The main rearrangements observed in our study are illustrated. Repetitive elements and genes involved in rearrangements are shown as arrows. Red indicates GCR cassette genes, gray represents *SPLTRA11* and *SPLTRA12* inverted repeats, and black denotes aldo/keto reductase genes. Their positions relative to the centromeres, telomeres, subtelomeric homologous sequences (yellow rectangles), and each other are indicated (not to scale). **(A)** GCRs in strains containing IR-LTRs. BIR involving the right arm of chromosome I is depicted. Other BIR events involving chromosomes II and III are not shown. Blue arrows are inverted duplication. **(B)** GCRs in strains with deleted IR-LTRs.

Fourth, the unstable nature of *SPLTRA*.*11* and *SPLTRA*.*12* was previously reported by Albrecht et al., who showed that they were involved in gene amplification (Albrecht et al., 2000). Gene amplification was scored by lithium acetate resistance, resulting from amplification of the *nhe1*/*sod2* gene located less than 300 bp from our GCR cassette. Interestingly, in most of the clones carrying an inverted duplication (5/6), chromosome healing occurred through telomere addition and only one exhibited a secondary rearrangement involving BIR. This contrasts with observations in *S. cerevisiae*, where foldback inversions are predominantly resolved through homology-mediated secondary rearrangements (Li et al., 2020).

Upon deletion of *SPLTRA11*-*SPLTRA12*, breakpoints were more heterogeneous, and no adjacent inverted duplications were detected, consistent with the absence of a strong IR driving DSB formation. Similar to budding yeast (Chen and Kolodner, 1999), the spectrum of GCRs included terminal chromosomal deletions accompanied by BIR-mediated translocations, interstitial deletions, or de novo telomere additions (Figure 4 and 5B). Notably, in the absence of *SPLTRA11*-*SPLTRA12*, the predominant event leading to GCR formation involves homeologous recombination between two NADP-dependent aldo/keto reductase genes located on different chromosome arms. Overall, the design of the GCR system enables scoring of rearrangements mediated by BIR events and provides a valuable framework for further exploring BIR-related mechanisms in fission yeast, which are less well characterized than those in budding yeast. Using our assay, we detected long tracts of BIR extending up to 600 kb. To our knowledge, this is the first report of such extensive DNA synthesis by BIR in fission yeast. BIR in fission yeast was formally demonstrated in response to a single-ended DSB, with synthesis tracts limited to 14 kb (Xu et al., 2025). The prevalence of BIR events in GCR recovery is consistent with previous observations: although only four spontaneous canavanine-resistant isolates were analyzed in a prior study, one resulted from a homology-driven nonreciprocal translocation between the left arms of chromosomes I and II (Lyu et al., 2024). Studies using minichromosome assays have reported that one class of rearrangement corresponds to isochromosome formation involving the repetitive centromeric sequences, with little evidence of BIR involvement (Nakamura et al., 2008b; Onaka et al., 2020; Pai et al., 2023). For example, Pai et al., ruled out the involvement of BIR events in the formation of GCRs even when only 3 kb of homology was provided (Pai et al., 2023). Expectedly, in the assay used in the Irie et al study, no BIR events were detected due to the lack of sequence homology with other chromosomal regions, making that assay well-suited for detecting GCRs mediated by very little or no homology (Irie et al., 2019). Here, the assay we developed enables to score spontaneous BIR events under physiological conditions. In contrast, recent work by Xu et al. showed the involvement of BIR in pericentromeric-induced GCR formation only in a context where heterochromatin is lost (Xu et al., 2026). This difference suggests that BIR may contribute to genome rearrangements during normal cellular state, highlighting a broader relevance of this pathway for genome stability.

While GCR assays are extensively used in *S. cerevisiae*, they are less commonly applied in *S. pombe*. As a model organism, *S. pombe* offers unique advantages. It is a fission yeast with chromosome structures and DNA repair pathways that more closely resemble those of higher eukaryotes than *S. cerevisiae*. Therefore, this assay will be very useful for identifying new factors that influence chromosomal integrity and for conducting comparative analyses with other eukaryotic models. To explore this, we measured GCR rates in mutants with known or unknown roles in genome stability pathways. The first gene we targeted is *nup132*. Several studies showed that DNA exposed to potentially damaging events such as DSBs, replication fork stalling or telomere erosion, relocates to the nuclear periphery, especially to the NPCs (reviewed in (Lamm et al., 2021; Su et al., 2015)). Nup132 in fission yeast was shown to be involved in promoting efficient recombination-mediated DNA synthesis upon fork stalling but dispensable for stalled fork relocation to the nuclear periphery (Kim and Forsburg, 2022; Kramarz et al., 2020; Schirmeisen et al., 2024). Deletion of *nup132* leads to sensitivity to replication stress-inducing agents (such as hydroxyurea) but not to DSB-inducing agents (such as bleomycin) (Kramarz et al., 2020). Therefore, it is plausible that faulty repair of spontaneous or IR-mediated replication-associated breaks is more readily channeled into GCRs in *nup132* mutants. This is in agreement with a finding in budding yeast where anchoring of stalled replication forks at tri-nucleotide repeats, another type of at-risk DNA sequences, to the NPC prevents GCRs (e.g. (Su et al., 2015; Zhang et al., 2006)).

Finally, beyond providing a complementary system to *S. cerevisiae* for comparative genomics and repair pathway analysis, our assay in fission yeast can be used to identify genes previously unknown for their involvement in the formation of genomic rearrangements. As a proof of concept, we demonstrate that the H3-H4 histone binding protein Djc9, which is not conserved in *S. cerevisiae*, plays a role in promoting GCR formation. Djc9 is the ortholog of human DNAJC9 and was recently found to promote histone H3-H4 degradation in fission yeast (Ding et al., 2025). Deletion of Djc9 allows to bypass the essentiality of the histone chaperone Asf1. It was suggested that, in the presence of Asf1, Djc9 may help remove surplus or defective histones and thus increase tolerance to replication stress and deleterious forms of histone H3. The fact that deletion of djc9 leads to a decrease in the GCR level suggests that increasing the pool of available wild-type histones plays a protective role against rearrangement formation. Interestingly, this role appears more prominent in the presence of a fragile motif, since deletion of *djc9* in the absence of *SPLTRA11-SPLTRA12* resulted in only a 1.5-fold decrease. The mechanism by which deletion of Djc9 mitigates GCR formation remains unknown, though. One possibility is that Djc9 promotes the initial event that triggers GCR formation (e.g., secondary structure formation), while another suggests a role in GCR recovery (e.g., by promoting telomere addition or BIR events). Both possibilities may relate to Djc9’s role in modulating histone availability (Ding et al., 2025). Further studies will be conducted to determine how Djc9 facilitates chromosome instability under physiological conditions.

In conclusion, we developed a convenient assay based on canavanine resistance to monitor GCR formation in fission yeast. It offers an opportunity to systematically examine how the deficiency of 356 *S. pombe*-specific genes, which are orthologs of many human disease genes (White and Allshire, 2008) affects the IR-induced or spontaneous GCR formation.

## Material and methods

### Plasmid and strain construction

The GCR cassette was constructed in several steps. In all PCR amplification reactions, *S. pombe* genomic DNA extracted from the HJ10 strain (h+ *leu1-32, ura4-D18*) was used as template. The chromosome I region I:50132-51288 (Pombase reference) was PCR-amplified using hybrid primers and inserted between the SalI and PspOMI sites on the pGEM5Zf(-) plasmid (Promega). The MunI site in the chromosome I region was converted into BamHI using annealed oligonucleotides. *ade7* was PCR-amplified from genomic DNA using hybrid primers containing two flanking BglII sites and one AscI site; the PCR product was digested with BglII, and the *ade7* fragment with AscI was inserted into BamHI. *cat1* was PCR amplified with hybrid primers containing two flanking MluI sites, and the digested fragment was inserted into the AscI site. Finally, *aat1* was PCR-amplified from genomic DNA using hybrid primers containing two flanking NsiI sites, and the aat1 NsiI fragment was inserted into the PstI site between the *ade7* and *cat1* genes. The resulting pKL718 plasmid was digested with SacI and the GCR cassette fragment flanked by homologous sequences to chromosome I was used to introduce the GCR cassette into the KT1937 strain (h+ *cat1Δ aat1Δ ade7Δ ura4-D18 leu1-32*), where *ade7, cat1, and aat1* were subsequently deleted using *delitto perfetto* (Fréon et al., 2023). This technique was also employed to delete the region encompassing *SPLTRA11* and *SPLTRA12*, and standard genetic methods were used to produce other deletion mutants. Primer design is available upon request. Canavanine sensitivity of arginine permease mutants was assessed by a spot test assay. Cultures from exponentially growing cells were serially 10-fold diluted in water and spotted on YES medium (control) or EMM medium containing 100 mg/L of canavanine (Sigma, C1625) and 0.5 g/L ammonium chloride, supplemented with adenine, leucine, and uracil (125 mg/L each).

### Fluctuation test to determine GCR rates

Cells were grown on YES plates and at least 11 individual colonies were selected for analysis. The individual colonies were resuspended in 200 µl of water and then serially diluted. Cells from appropriate dilutions were plated on YES media for survival and canavanine-containing EMM media supplemented with low amount of adenine (100 mg/L canavanine, 0.5 g/L ammonium chloride, 12.5 mg/L adenine, 125 mg/L uracil, 125 mg/L leucine). After 5-10 days of incubation at 30°C, colonies were counted to determine the GCR rate. Red colonies were counted on canavanine-containing media (Can^R^ Ade^-^). The GCR rate and confidence intervals were determined as described in (Drake, 1991), using the formula μ = f/ln(Nμ). Data are represented as the median rate ± 95% confidence interval.

### Whole genome sequencing

Genomic DNA was extracted from the GCR clones (Can^R^Ade^-^) using the YeaStar Genomic DNA kit (Zymo research, D2002) with the following modification: after zymolyase treatment, the spheroplast suspension was incubated for 60 minutes with 10 ul of protease K (10mg/ml) at 50Co before it was applied to the column. Illumina sequencing (2x150 bp) was performed by Azenta Life Science, with ∼1 GB of coverage per sample. Reads were mapped to the reference sequence ASM294v2.62 using QIAGEN CLC Genomics Workbench.

### Rearrangement and read-depth analysis

Chromosome rearrangements were identified using the CLC Genomics Workbench software. Chromosomal read coverage computation and plotting were performed using an in-house Python script (using pysam and numpy libraries). Briefly, per-base coverage was computed using the pileup method followed by Reads Per Million (RPM) normalization to facilitate multiple samples comparisons. The normalized signal was finally averaged over 1 kb binned genome. The Y-axis was scaled to 25RPM for visual comparison ease.

### Inverted dimer detection by Southern blot hybridization

Genomic DNA was extracted from 1x10^8^ cells grown in YES medium. DNA was RNase-treated and submitted to restriction digestion with 15 units of NsiI (NEB R0127S). DNA fragments were resolved by electrophoresis in 0.8% agarose gel in 1x TBE. DNA was then transferred to a GeneScreen Plus nylon membrane (Perkin-Elmer, NEF988001PK) by capillary in 10x SSC. The membrane was probed with a ^32^P-radiolabeled (TaKaRa, 6046) DNA fragment and incubated in Ultra-Hyb buffer (Invitrogen, AM8669) at 42°C overnight. The probe used corresponds to a sequence located downstream of *SPLTRA12*. After washing, the membrane was exposed to a phosphor storage screen, and the signal was collected using the phosphor imager software on a Typhoon Trio.

### Data access

All raw and processed sequencing data generated in this study have been submitted to the NCBI Gene Expression Omnibus (GEO; https://www.ncbi.nlm.nih.gov/geo/) under accession number GSE327142.

## Author contributions

A.A.S., C.O., A.B.C. and K.S.L. performed the experiments. A.A.S. and K.S.L. contributed to experimental design and data analysis. K.M. provided bioinformatic expertise for the depth read analysis. S.A.E.L. provided scientific discussion and expertise. A.A.S. and K.S.L. wrote the manuscript. All authors reviewed and edited the manuscript.

