## Supplemental data for "Canavanine-based assay for gross chromosomal rearrangements reveals genome instability hotspots and modulating genes in fission yeast"

**Table S1.** GCR rates in *nup133Δ* in *S. cerevisiae*.

| Strain | Rate (CI95%) | Fold change (relative to WT) |
|---|---|---|
| WT | $6.82x10^{-7}$ ($5.29x10^{-7}$-$1.12x10^{-6}$) | |
| *nup133Δ* | $3.82x10^{-7}$ ($2.83x10^{-7}$-$4.89x10^{-7}$) | 1.8 ↓ |

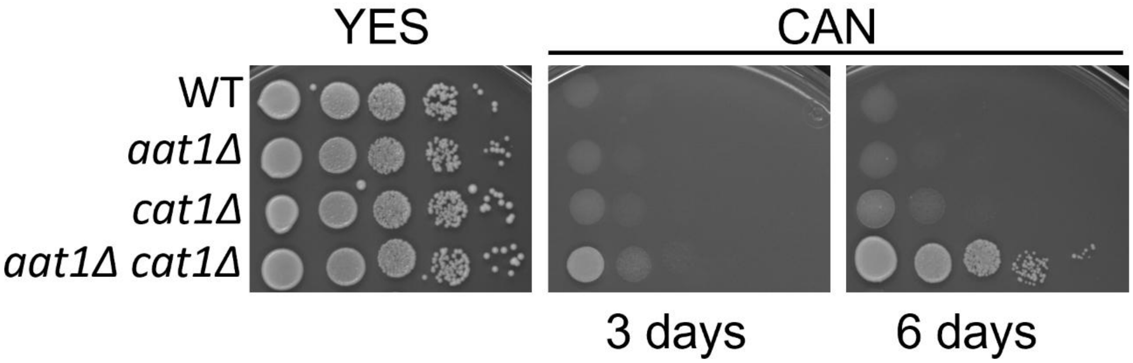


**Figure S1.** Deletions of *cat1* and *aat1* have additive effects on canavanine resistance. Canavanine resistance in indicated strains was assayed by serial dilutions on EMM media containing 0.5 g/L of ammonium chloride and 100 mg/L of canavanine.

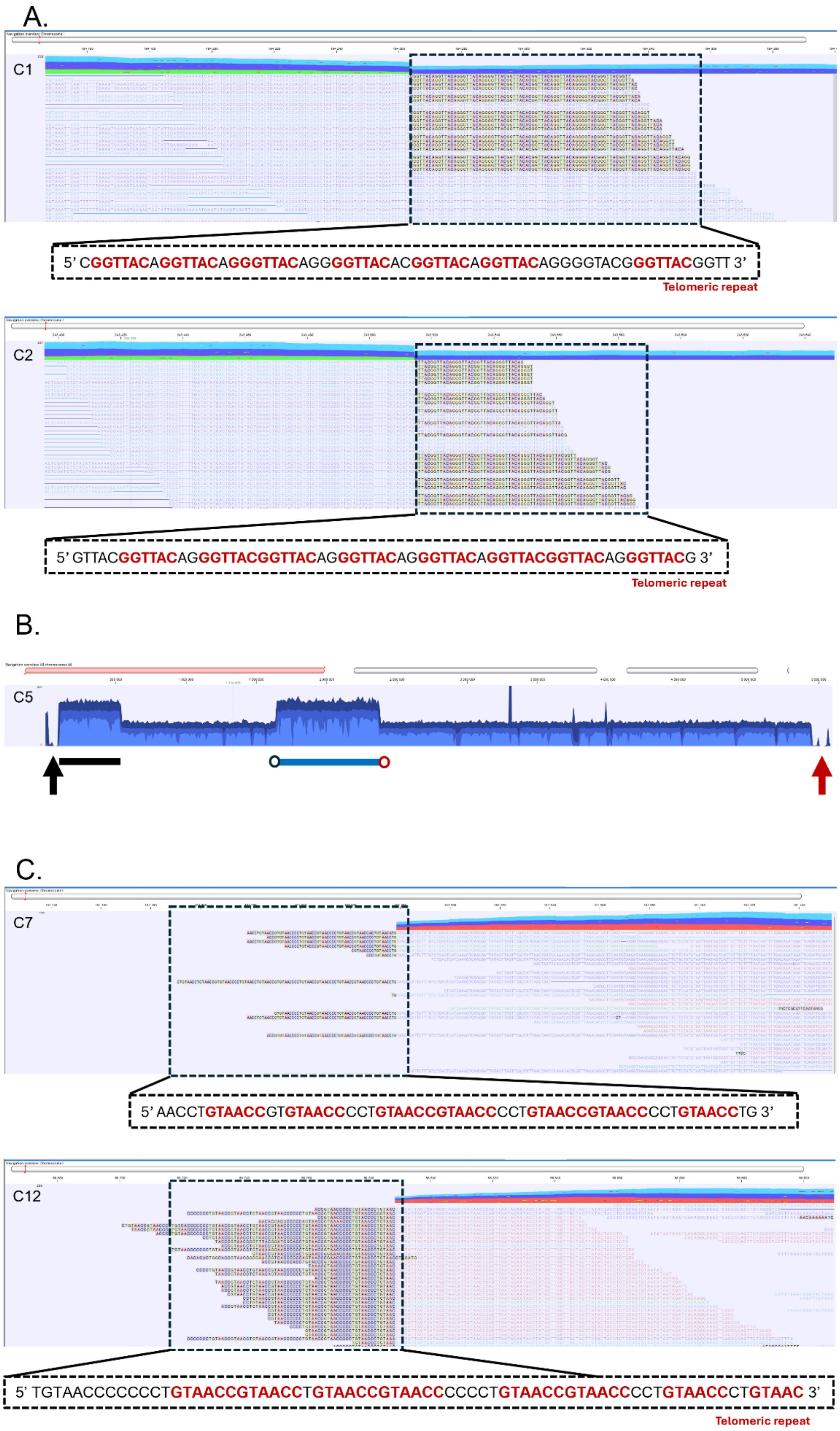
A.
C1
5' CGGTTACAGGTTACAGGGTTACAGGGGTTACACGGTTACAGGTTACAGGGGTACGGGTTACGGTT 3'
Telomeric repeat
C2
5' GTTACGGTTACAGGGTTACGGTTACAGGGTTACAGGGTTACAGGTTACGGTTACAGGGTTACG 3'
Telomeric repeat
B.
C5
C.
C7
5' AACCTGTAACCGTGTAACCCCTGTAACCGTAACCCCTGTAACCGTAACCCCTGTAACCTG 3'
C12
5' TGTAACCCCCCCTGTAACCGTAACCTGTAACCGTAACCCCCCTGTAACCGTAACCCCTGTAACCCTGTAAC 3'
Telomeric repeat

**Figure S2.** Examples of GCR events resolved by telomere addition. **A.** Examples of telomere addition in clones showing an inverted duplication of chromosome I. Unmapped sequences at the duplication end correspond to telomeric repeats. B. global view of chromosome I from clone C5. Black arrow: terminal deletion of the left arm. Red arrow: terminal deletion of the right arm. Black line: adjacent duplication from the left breakpoint. Blue line: secondary duplication. Black circle: position of unmapped sequences corresponding the end of the adjacent duplication (telomere distal). Red circle: position of unmapped sequences corresponding to the right arm breakpoint region. **C.** Detection of telomeric repeats adjacent to the breakpoint in GCR clones devoid of secondary rearrangements.

**Table S2.** Strains used in this study.

| Name | Genotype* |
|---|---|
| AA823 | *h+ cat-DP** ade7-DP aat1-DP ura4-D18 leu1-32* |
| AA997 | *h-smt0 ChrI-leftarm:ade7-cat1-aat1 cat1-DP ade7-DP aat1-DP ura4-D18 leu1-32* |
| AA1046 | *h+ SPLTRA.11-SPLTRA.12-DP ChrI-leftarm:ade7-cat1-aat1 cat1-DP ade7-DP aat1-DP ura4-D18 leu1-32* |
| AA1072 | *h+ nup132::NAT ChrI-leftarm:ade7-cat1-aat1 cat1-DP ade7-DP aat1-DP ura4-D18 leu1-32* |
| AA1082 | *h+ nup132::NAT SPLTRA.11-SPLTRA.12-DP ChrI-leftarm:ade7-cat1-aat1 cat1-DP ade7-DP aat1-DP ura4-D18 leu1-32* |
| AA1117 | *h+ djc9::Nat ChrI-leftarm:ade7-cat1-aat1 cat1-DP ade7-DP aat1-DP ura4-D18 leu1-32* |
| AA1147 | *h+ djc9::natMX SPLTRA.11-SPLTRA.12-DP ChrI-leftarm:ade7-cat1-aat1 cat1-DP ade7-DP aat1-DP ura4-D18 leu1-32* |
| AA1154 | *h+ aat1-DP ura4-D18 leu1-32* |
| AA1156 | *h+ cat-DP ade7-DP ura4-D18 leu1-32* |

*All strains have been generated in this study

** DP for deletion by *delitto perfetto.*
